# Requirement of hypoxia-inducible factor 1 alpha for interleukin 1 beta induced glycolysis in colorectal cancer cells

**DOI:** 10.64898/2026.08.11.744327

**Authors:** Ji Yeon Kim, Bohye Park, Olivia F. Riffey, Ahmed Bettaieb, Dallas R. Donohoe

## Abstract

Colorectal cancer cells increase glycolysis to help meet the metabolic demands required for cell growth. Many factors, both endogenous and exogenous, likely drive cellular metabolism and enhance glycolytic flux in colorectal cells. Interleukin-1 beta (IL-1β) is a pro-inflammatory cytokine that is elevated in colorectal cancer. In this study, we investigated the effect of IL-1β toward driving the cancer cell to increase glycolysis, while also suppressing the oxidation of the fiber-derived nutrient butyrate. The results presented here demonstrate that IL-1β stimulated glycolysis and inhibited maximal mitochondrial respiration. IL-1β also increased the phosphorylation of AKT and hypoxia-inducible factor 1 alpha (HIF1α) levels. Utilizing colorectal cancer cells with AKT1/2 or HIF1α knocked out showed the requirement of these proteins in mediating the increase in glycolysis following IL-1β treatment. Importantly, AKT1/2 was identified as upstream of HIF1α, as IL-1β still increased phosphorylation of AKT even in the absence of HIF1α. However, loss of AKT1/2 completely abolished the ability of IL-1β to increase HIF1α protein levels. Tumor necrosis factor alpha (TNFα), another cytokine found to be elevated in colorectal cancer, also increased glycolysis in an AKT and HIF1α-dependent manner. Our data point to a common pathway through AKT activation and HIF1α upregulation, by which pro-inflammatory cytokines increase glycolysis in colorectal cancer cells to help promote cancer progression.

## Introduction

Over the past several years, cellular metabolism in the colonic epithelial cell or colonocyte has been highlighted as a major contributor in regulating microbiome composition, including pathogen susceptibility and gastrointestinal health and disease [1–4]. A central player in colonocyte metabolism is the oxidation of the short-chain fatty acid butyrate, which serves as the primary energetic substrate for non-cancerous colonic epithelial cells [5, 6]. Reductions in the availability or oxidation of butyrate in the colonic epithelium has been associated with reduced barrier function, a decrease in the number of colonic stem cells, and enhanced luminal oxygen and nitrogen levels in the colon [1, 7–9]. At the same time, reduced butyrate oxidation has been associated with increased glycolysis in colonocytes [3]. Cancerous colonocytes show elevated glycolysis, which is a key feature in the progression of colorectal cancer [10]. Increases in glycolysis may involve an upregulation of the AKT/PI3K-signaling pathway in cells [11]. Upregulation of the AKT/PI3K pathway increases fructose-2, 6-biphosphate through activation of 6-phosphofructo-2-kinase/fructo-2, 6-biphosphatase (PFKFB2) [12]. An elevation in fructose-2, 6-biphosphate allosterically activates the glycolytic enzyme phosphofructokinase 1 (PFK1) to increase glucose flux or glycolysis [13]. Another mediator of increased glycolysis in cells is a hypoxia-inducible factor (HIF1α), which regulates the transcription of glycolytic enzymes, leading to promoting glycolysis and supporting tumor survival and progression [14–17]. In addition to increasing glycolysis, cancerous colonocytes or colorectal cancer cells show a diminishment in butyrate oxidation [3, 18].

Several factors likely promote this metabolic transition towards increased glycolytic flux, including the environmental niche surrounding the colorectal cancer. Elevated proinflammatory cytokines, such as interleukin-1 beta (IL-1β) and tumor necrosis factor alpha (TNFα), are components of this niche and are able to directly act on colorectal cancer cells and primary colon carcinoma organoids [19–21]. IL-1β promotes glycolysis in lung adenocarcinoma cells, macrophages, colorectal cancer cells, and osteoarthritic chondrocytes [19, 22–24]. However, it is not entirely clear how IL-1β would affect glycolysis in colorectal cells.

Colorectal cancer (CRC) can sometimes take years to manifest as the colorectal tumor (adenoma) progresses into an invasive carcinoma. Inflammation and diet are two factors that interact to affect this process [25–27]. In the case of inflammation, elevated pro-inflammatory cytokines such as TNFα and IL-1β increase colitis-associated and non-colitis-associated colorectal cancer [26, 28, 29]. In contrast to the pro-inflammatory cytokines, dietary factors such as fiber are suggested to lower CRC susceptibility [30]. The interplay between inflammatory factors and dietary factors together also plays a role in regulating tumor progression. Importantly, although much is known as to how diet impacts inflammation, far less is understood about how inflammatory mediators impact nutrients and their metabolism. Here, we demonstrate that IL-1β and TNFα at physiologically-relevant levels, induce a metabolic shift in colorectal cancer cells towards increased glycolytic flux, while inhibiting the oxidation of the fiber metabolite butyrate. Moreover, we find that AKT and HIF1α are necessary for this metabolic shift. These data allude to the mechanisms contributing to the metabolic shift that occurs in colorectal cancer cells, which is partly promoted through the pro-inflammatory cytokines IL-1β and TNFα.

## Materials and methods

### Cell Culture

HCT 116 colorectal cancer cells were purchased from the American Type Culture Collection (ATCC, #CCL-247). HCT 116 HIF1α knockout cells and HCT 116 AKT1 and AKT2 (AKT1/2) double knockout cells were provided by Dr. Bert Vogelstein (Johns Hopkins University School of Medicine). They were originally generated using an adeno-associated virus (AAV)-mediated homologous recombination for HIF1α knockout cells and a recombinant AAV-based promoter-trap strategy for AKT1/2 KO cells [31, 32]. All cells were maintained in DMEM supplemented with 25 mM glucose, 10% fetal bovine serum (FBS), and 1% penicillin/streptomycin at 37 °C in 5% CO2 environment and passaged following the methods described in earlier studies [33]. In brief, cells were passaged by aspirating the culture medium, washing with PBS, and treating with 0.25% Trypsin-EDTA until detached. Trypsin was inactivated with culture medium, cells were dispersed by pipetting, and reseeded at a split ration of 1:5 to 1:7. For treatment, the cells were grown to 70% confluence and treated with human recombinant IL-1β or TNFα (PeproTech, Inc., Rocky Hill, CT, USA) for 6 h in DMEM supplemented with 5 mM glucose, and 10% FBS.

### Flux experiments

The Seahorse XFe24 Analyzer (Agilent Technologies, CA) was used for real-time measurements of glycolytic proton efflux rate (glycoPER), oxygen consumption rate (OCR), and extracellular acidification rate (ECAR). The glycolytic rate assay, mito stress assay, and glycolysis stress test were performed according to the manufacturer’s instructions. Cells were seeded at a density of 2.5 × 10^5^ cells/well 24 hours before the assay. Six hours before the assay, cells were treated with 1 ng/mL IL-1β or TNFα. One hour before measuring glycoPER, OCR or ECAR, cells were maintained with XF Assay medium (Agilent, 103575) for glycolytic rate assay and mito stress assay, and XF Assay medium (Agilent, 102353-100) for glycolysis stress test. In the glycolytic rate assay, analysis was conducted in the XF Assay medium following the addition of Rotenone/Antimycin (0.5 μM) and 2-deoxy-glucose (2DG) (50 mM). In the mito stress assay, analysis was performed in the XF Assay media following the addition of oligomycin (1 μM), Carbonyl cyanide-4 (trifluoromethoxy) phenylhydrazone (FCCP) (1 μM), and Rotenone and Antimycin (1 μM). In the glycolysis stress test, analysis was conducted in the XF Assay media following the addition of glucose (10 mM), oligomycin (1 μM), and 2DG (50 mM).

The butyrate oxidation assay was optimized in our previous work [18]. Briefly, cells were seeded in an XFe24 cell plate at 2.5 × 10^5^ cells/well. 6 hours before the assay, cells were treated with IL-1β. One hour before the measurement of butyrate oxidation, the medium was changed to 1X Krebs-Henseleit Buffer (KHB), including 5 mM glucose and 500 μM carnitine, and the cell plate was incubated in a non-CO2 incubator at 37 °C for one hour. The butyrate oxidation assay was performed using sequential injections of 5 mM sodium butyrate, 50 mM 2DG, and 10% sodium azide (SA). After each assay, cells were harvested directly on the XF cell plate with 1X RIPA buffer, and the lysates were quantified using a Pierce BCA Protein Assay kit (Thermo Fisher, PI23228) to normalize each assay.

### Glucose uptake assay

Glucose uptake was quantified using a glucose uptake assay obtained from Promega (Madison, WI, USA). HCT 116 colorectal cancer cells were seeded at a density of 2.5 × 10^5^ cells/well in 96-well plates one day before the assay. Six hours before the assay, IL-1β (1 ng/mL) stimulated HCT 116 cells. Before beginning the assay, the medium—including IL-1β—was removed, and the cells were washed with 100 μl of phosphate-buffered saline (PBS). To initiate glucose uptake, 50 μl of 2DG (1 mM) in PBS was added to the cells. The uptake reaction was stopped, and the samples were processed as described in the manufacturer’s instructions. All data were acquired on a multi-well plate reader (BioTek Synergy HTX multi-mode plate reader, USA) with an integration time of 0.5 s and normalized to total cell protein measured by Pierce BCA protein assay kit.

### Western blot analysis

Cells were seeded at a density of 2 × 10^6^ cells in 100 mm cell culture dish and the cells were grown to 70% confluence and treated with 1 ng/ml of IL-1β or TNFα for 6 h in DMEM supplemented with 5 mM glucose, and 10% FBS. After IL-1β or TNFα treatment, cells were washed with twice with 1X PBS and lysed in lysis buffer made with 1X RIPA buffer (Cell Signaling, 9806s), 1 mM PMSF (Cell Signaling, 8553), 0.5 M EDTA (Thermo Scientific, R1021), and HaltTM phosphatase inhibitor cocktail (Thermo Scientific, PI87786). Protein concentrations were determined with the BCA assay kit (Thermo Scientific, 23227). The proteins (20 µg) were separated by 8 or 10% SDS-polyacrylamide gels and transferred onto PVDF membranes. After being blocked with 5% non-fat milk or 5% BSA in 1X TBST (0.1%) for 1 h at room temperature, the membranes were incubated with AKT, phospho-AKT, HIF1α, or β-actin (Cell Signaling, 3700; 1:5000) overnight at 4 °C. The membranes were washed three times with 1X TBST (0.1%) for 10 min each time at room temperature. They were then incubated for 1 h at room temperature with either fluorescent secondary antibodies (LI-COR, #50489565; Licor, #92568022) diluted 1:10000 or horseradish peroxidase-conjugated secondary antibodies (BOSTER, #BA1054; Novus, #NB7570) diluted 1:10000 for 1 h at room temperature. This was followed by washing the membranes three times in 1X TBST (0.1%) for 10 min each time. Fluorescent or chemifluorescent signals of the membrane were detected using Odyssey Fc (LI-COR Biosciences, Lincoln, NE, USA), and Image Studio software (LI-COR, Biosciences, Lincoln, NE, USA) was used for the quantification of the signal intensities.

### Antibody List

Table 1 provides the Western blot antibody list.

**Table 1.** Western Blot Antibody List.

| **Antibody target (origin)** | **Dilution** | **Company** | **Catalog number** |
| --- | --- | --- | --- |
| AKT (Rabbit) | 1:1000 | Cell Signaling | 4691s |
| phospho-AKT (Rabbit) | 1:1000 | Cell Signaling | 4060s |
| HIF1α (Rabbit) | 1:500 | Cell Signaling | 3716s |
| β-actin (Mouse) | 1:5000 | Cell Signaling | 3700 |
| Goat anti-Rabbit IgG<br>Secondary Antibody (Rabbit) | 1:10000 | Li-cor | 926-32211 |
| Donkey anti-Mouse IgG<br>Secondary Antibody (Mouse) | 1:10000 | Li-cor | 926-68022 |
| HRP Conjugated Goat Anti-Rabbit IgG (H+L) secondary<br>Antibody (Rabbit) | 1:10000 | Boster | BA1054 |
| HRP Conjugated Goat anti-Mouse IgG (H+L) Secondary<br>Antibody | 1:10000 | Novus | NB7570 |

### Statistical analysis

For seahorse experiments, glucose uptake assays, and western blot, t-test or one-way ANOVA was used to test for differences between experimental groups, followed by a Tukey post-hoc test for the one-way ANOVA. All data are expressed as mean ± SEM. Significant differences are indicated.

## Results

### IL-1β increases glycolysis while diminishing butyrate oxidation in colorectal cells

Utilizing the Seahorse XF Analyzer, we found that the colorectal cancer cells that were incubated for at least 6 h with 1 ng/mL IL-1β showed a significant increase in their extracellular acidification rate (ECAR), which is indicative of lactate and carbon dioxide excretion by the cells (**Fig S1A&B**). These data showed that chronic (6 h exposure), but not acute IL-1β exposure promoted significant effects on colorectal cell metabolism. However, the exact glycolytic contribution could not be separated from the carbon dioxide component of the ECAR measurement.

To test the glycolytic component of the ECAR, rotenone and antimycin A were used to block mitochondrial respiration and carbon dioxide production in a glycolytic rate assay. The mitochondrial proton efflux rate is subtracted from the total proton efflux rate to achieve the glycolytic proton efflux rate or glycoPER. Treatment of colorectal cancer cells for 6 h significantly elevated the glycoPER (**Fig 1A**). This was judged by the increased basal (**Fig 1B**) and compensatory glycolysis (**Fig 1C**). The basal glycolysis represented the first three measurements of glycoPER. The compensatory glycolysis corresponds to the measurements after the addition of rotenone and antimycin A, but before the injection of 2-deoxyglucose (2DG).

**Figure 1.**
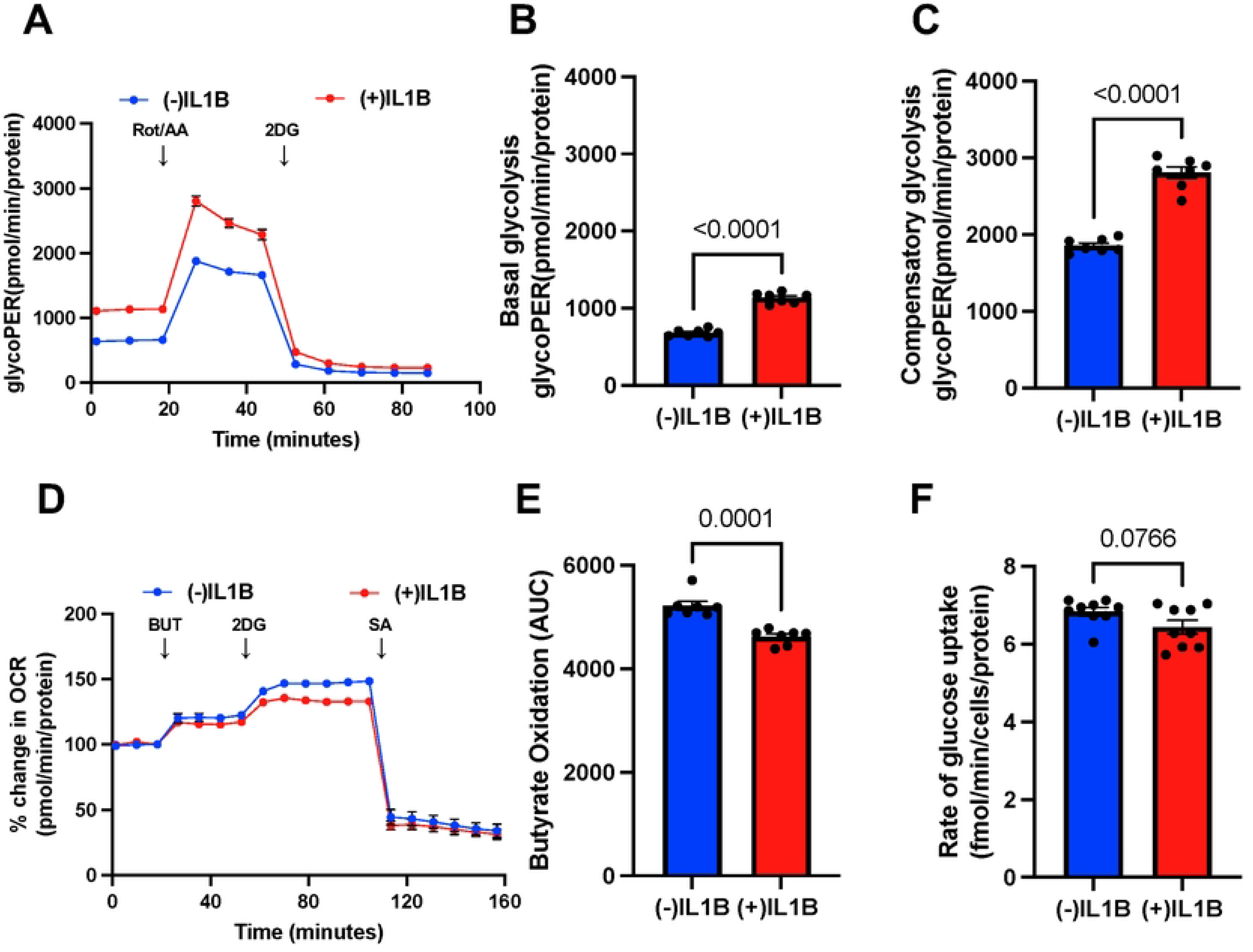
Interleukin-1 beta induces glycolysis while suppressing butyrate oxidation. Contribution of glycolysis to proton efflux rate (glycoPER). (**A**), basal glycolysis (**B**), and compensatory glycolysis (**C**) in untreated control and IL-1β-treated HCT 116 colorectal cancer cells. Oxygen consumption rate (OCR) relative to baseline in untreated and IL-1β-treated HCT 116 cells with and without butyrate (5 mM) (**D**). The total contribution of butyrate toward OCR is observed after injection of 2-deoxyglucose (2DG). Butyrate oxidation (**E**) represents the area under the curve (AUC) analysis from OCR measurements taken after 2DG injection but before sodium azide (SA) injection. Data points represent the average OCR over 4–5 replicates per condition for butyrate oxidation measurements. The rate of glucose uptake in untreated and IL-1β-treated HCT 116 cells (**F**). Error bars are mean ± SEM and ns indicates no statistical significance (P > 0.05). Significant differences are shown.

Treatment of colorectal cancer cells with IL-1β also resulted in diminished butyrate oxidation (**Fig 1D**). Butyrate oxidation in colorectal cells was determined after the addition of 2DG, where butyrate is the only exogenous carbon source (**Fig 1E**). IL-1β showed no effects on glucose uptake (**Fig 1F**). The effect of IL-1β toward diminishing butyrate oxidation was observed after only 45 min (**Fig S2**). Taken together, these data suggest that IL-1β elicits a shift in cellular metabolism away from butyrate oxidation and towards glycolysis and glucose consumption.

### IL-1β inhibits mitochondrial respiratory capacity

Based on the reduced butyrate oxidation caused by IL-1β, we decided to interrogate mitochondrial function. Treatment of colorectal cancer cells with IL-1β but not butyrate did not alter the basal oxygen consumption rate (OCR) (**Fig 2A**). However, IL-1β did lower the maximal respiratory capacity (**Fig 2B**) and spare respiratory capacity (**Fig 2C**). Interestingly, there was a small, but significant, increase in the non-mitochondrial oxygen consumption (**Fig 2D**). IL-1β treatment of colorectal cancer cells showed no effect on the ATP production-coupled respiration (**Fig 2E**). Thus, although IL-1β treatment increased glycolysis and diminished the oxidation of butyrate, it did not drastically alter mitochondrial function. This suggests that the effect of IL-1β is selective for butyrate oxidation and not glucose. To explore the underlying mechanism, we considered known IL-1β signaling pathways. IL-1β binds to the interleukin-1 receptor where it activates several downstream proteins, including AKT/PI-3K [34]. AKT/PI-3K increases glycolytic flux through phosphorylating phosphofructokinase-2 and elevating fructose 2,6-biphosphate levels, which allosterically activates phosphofructokinase-1 (PFK-1) to help drive glycolysis [13].

**Figure 2.**
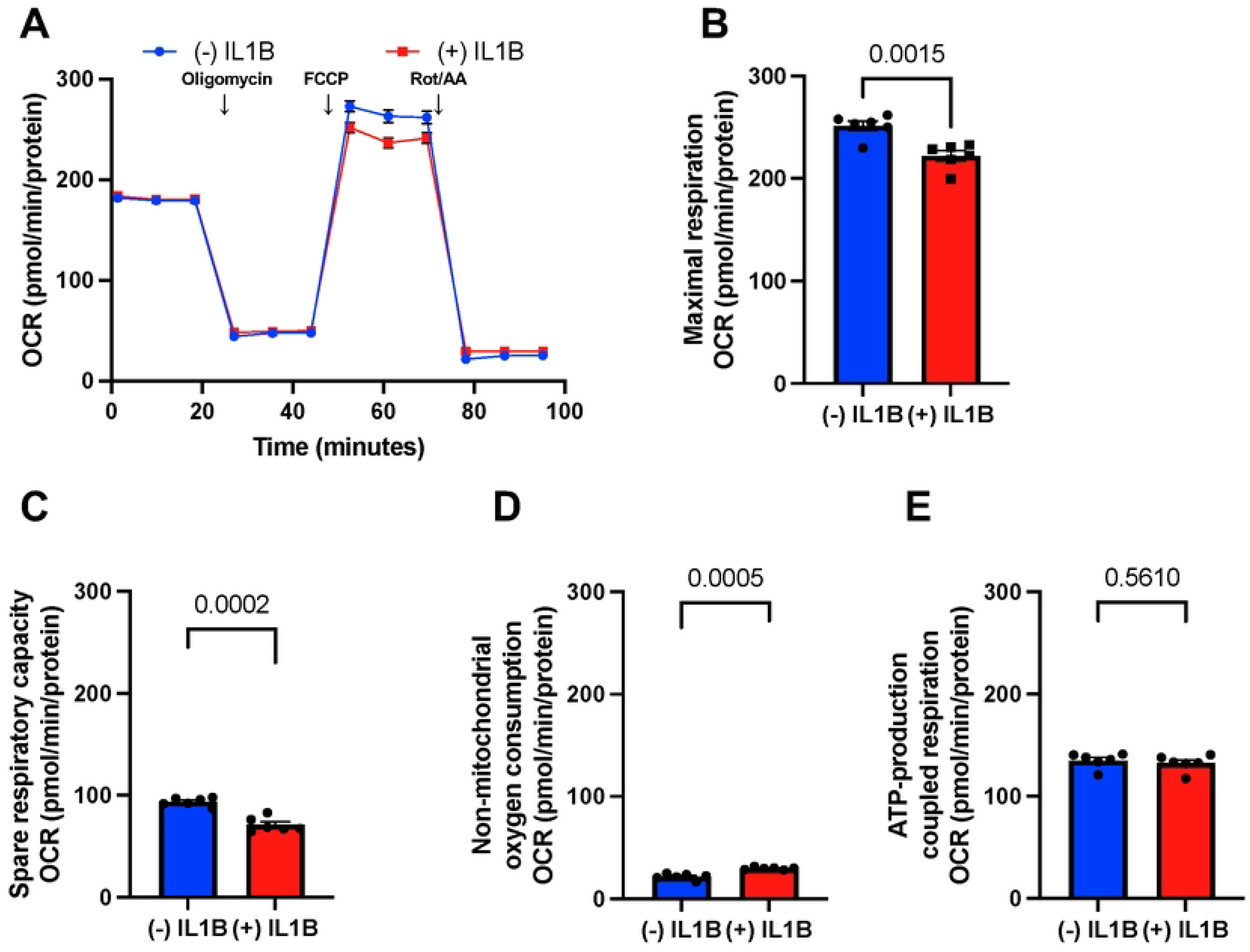
Mitochondrial function is inhibited by interleukin-1β. (**A**) Oxygen consumption rate (OCR) relative to baseline, in which untreated and IL-1β-treated HCT 116 cells following response to oligomycin, FCCP, and antimycin A/rotenone. The calculated maximal respiration (**B**), spare respiratory capacity (**C**), non-mitochondrial oxygen consumption (**D**), and ATP-coupled respiration (**E**). These measurements represent the mitochondrial function. Each data point represents the average OCR over 3–5 replicates per condition. Error bars are mean ± SEM and ns indicates no statistical significance (P > 0.05). Significant differences are shown.

### IL-1β increases Hypoxia Inducible Factor-1 Alpha through AKT

Hypoxia inducible factor is a transcription factor that regulates several glycolytic enzymes, including lactate dehydrogenase A (LDHA), pyruvate dehydrogenase kinase-1 (PDK-1), pyruvate kinase M2 (PKM2), and glucose transporter-1 (GLUT1) [3, 35]. An elevation in HIF1α is associated with an upregulation of these enzymes and increased glycolysis [36]. IL-1β has been previously shown to increase HIF1α levels in a human lung epithelial cell line [37]. HIF1α would represent a plausible mechanism as to how IL-1β induces glycolysis in HCT 116 cells. Towards this end, treatment of HCT 116 colorectal cancer cells with IL-1β did indeed significantly increase HIF1α levels (**Fig 3A**). This increase in HIF1α was also found to be dependent on AKT1/AKT2, as HCT 116 cells lacking these two enzymes did not show the same response to IL-1β (**Fig 3B**). Thus, these data show that IL-1β increases HIF1α through AKT1/AKT2 in colorectal cancer cells.

**Figure 3.**
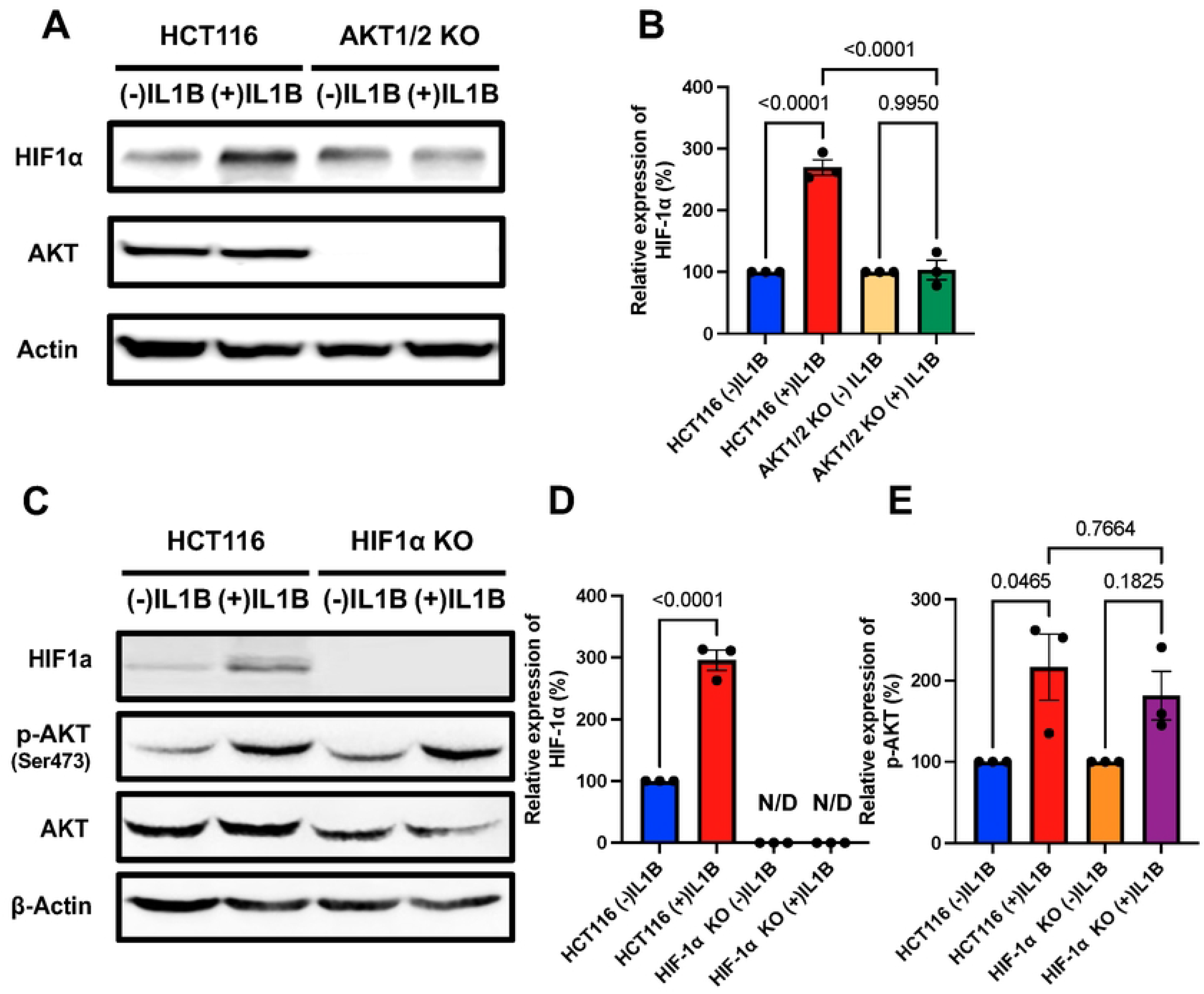
Interleukin-1β increases HIF1α through AKT activation in colorectal cancer cells. (**A**) Western blot analysis of HIF1α and AKT expressions in IL-1β-treated or untreated HCT 116 cells, or HCT 116 AKT1/2 knockout cells with β-actin as a loading control. (**B**) Quantification of the western blot is shown. (**C**) Western blot analysis of HIF1α, phospho-AKT (Ser473), and total AKT expression in IL-1β-treated or untreated HCT 116, or HCT 116 HIF1α knockout cells with β-actin as a loading control. Quantification of HIF1α (**D**) and phospho-AKT relative to total AKT (**E**). For statistical analysis, western blots were conducted at least 3 times. Error bars are mean ± SEM and N/D indicates not dectected. Significant differences are shown.

Although we found that AKT1/AKT2 is upstream of HIF1α, it has also been shown that AKT1/AKT2 is downstream of HIF1α. HCT 116 cells treated with IL-1β displayed elevated phospho-AKT at Ser473 (**Fig 3C**) and increased HIF1α (**Fig 3D**). However, IL-1β was not able to significantly effect phospho-AKT in HCT 116 cells where HIF1α was knocked out (**Fig 3E**). These data suggest that HIF1α is required for phosphorylation of AKT (Ser473) induced by IL-1β.

### AKT and HIF1a mediate IL-1β induced elevation in glycolysis

To test whether the elevation in glycolysis observed when treating colorectal cancer cells with IL-1β was dependent on AKT, a glycolytic rate assay was performed with HCT 116 colorectal cancer cells with and without AKT1/AKT2 to observe whether IL-1β could increase glycolysis. Loss of AKT1/AKT2 completely abolished the ability of IL-1β to increase glycolysis in these cells (**Fig 4A**). Neither basal (**Fig 4B**) or compensatory (**Fig 4C**) glycolysis was affected by IL-1β in AKT1/2 knockout cells. As expected, the basal glycolysis in the AKT1/2 knockout cells was diminished compared to HCT 116 parental cells, 511 vs. 705 pmol/min/protein (**Fig 4B**). However, the fact these cells failed to respond to IL-1β in regards to increasing glycolysis demonstrates the requirement of AKT signaling in the effect. We also report that IL-1β had no effect on the oxygen consumption rate (OCR) in AKT1/2 knockout cells (**Fig 4D**). Moreover, the mitochondrial oxygen consumption rate (mitoOCR) relative to the glycolytic proton efflux rate (glycoPER) showed no significant difference in AKT1/2 cells treated with or without IL-1β (**Fig 4E**).

**Figure 4.**
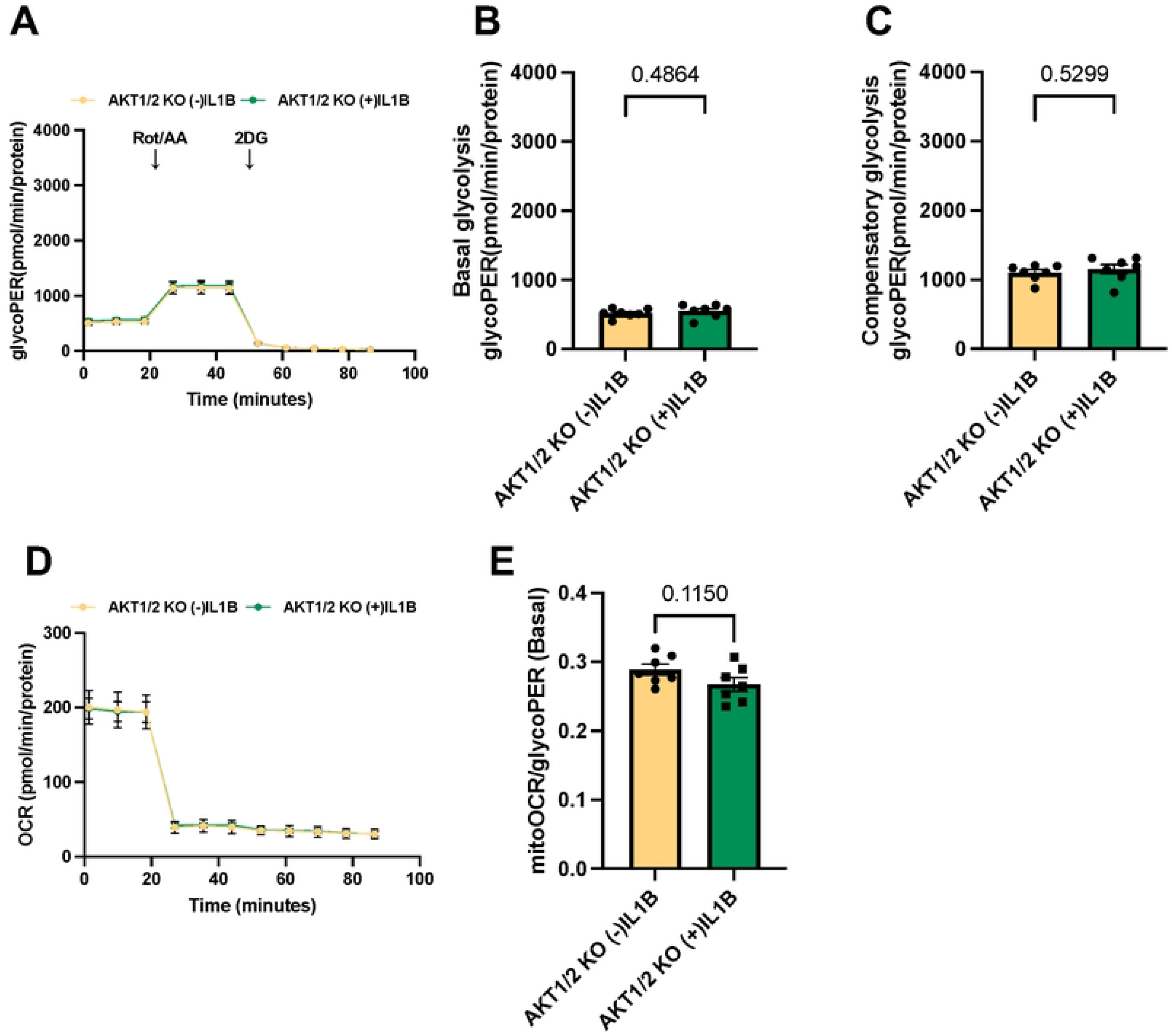
AKT knockout blocks interleukin-1β-induced glycolysis. Contribution of glycolysis to proton efflux rate (glycoPER) (**A**), basal glycolysis (**B**), and compensatory glycolysis (**C**) in untreated and IL-1β-treated HCT 116 AKT1/2 knockout colorectal cancer cells. Oxygen consumption rate (OCR) relative to baseline in untreated and IL-1β-treated HCT 116 AKT1/2 knockout cells (**D**). The basal mitochondrial oxygen consumption relative to the proton efflux from glycolysis (glycoPER) in HCT 116 AKT1/2 knockout cells treated with or without IL-1β (**E**). Each data point represents the average OCR over 3–5 replicates per condition. Error bars are mean ± SEM and ns indicates no statistical significance (P > 0.05)

We previously showed that loss of HIF1α completely abolished the ability of IL-1β to increase phosphorylation and activation of AKT, suggesting that HIF1α is the downstream target and mediator of increased glycolysis induced by IL-1β. Similar to the AKT1/2 knockout cells, loss of HIF1α eliminated IL-1β’s induction of glycolysis (**Fig 5A**). In addition, the glycoPER was lower in cells where HIF1α is knocked out (**Fig 5A**) as opposed to cells containing HIF1α (**Fig 1A**), confirming the importance of this transcription factor in promoting glycolysis. Basal (**Fig 5B**) and compensatory (**Fig 5C**) glycolysis also showed no difference. Furthermore, IL-1β no longer produced effects on mitochondrial function in HIF1α knockout cells (**Fig 5D**). IL-1β-treated HIF1α knockout cells exhibited no difference in maximal respiratory capacity (**Fig 5E**), spare respiratory capacity (**Fig 5F**), non-mitochondrial oxygen consumption (**Fig 5G**), and ATP-coupled respiration (**Fig 5H**) compared to non-treated cells. Taken together, these data demonstrate that IL-1β increases glycolysis through phosphorylation of AKT, which then functions through HIF1α. In addition, loss of either AKT1/2 or HIF1α is completely sufficient to mediate IL-1β’s effects on glycolysis. AKT and HIF1α represent common targets, and as such, additional cytokines have been previously shown to function through these proteins [38].

**Figure 5.**
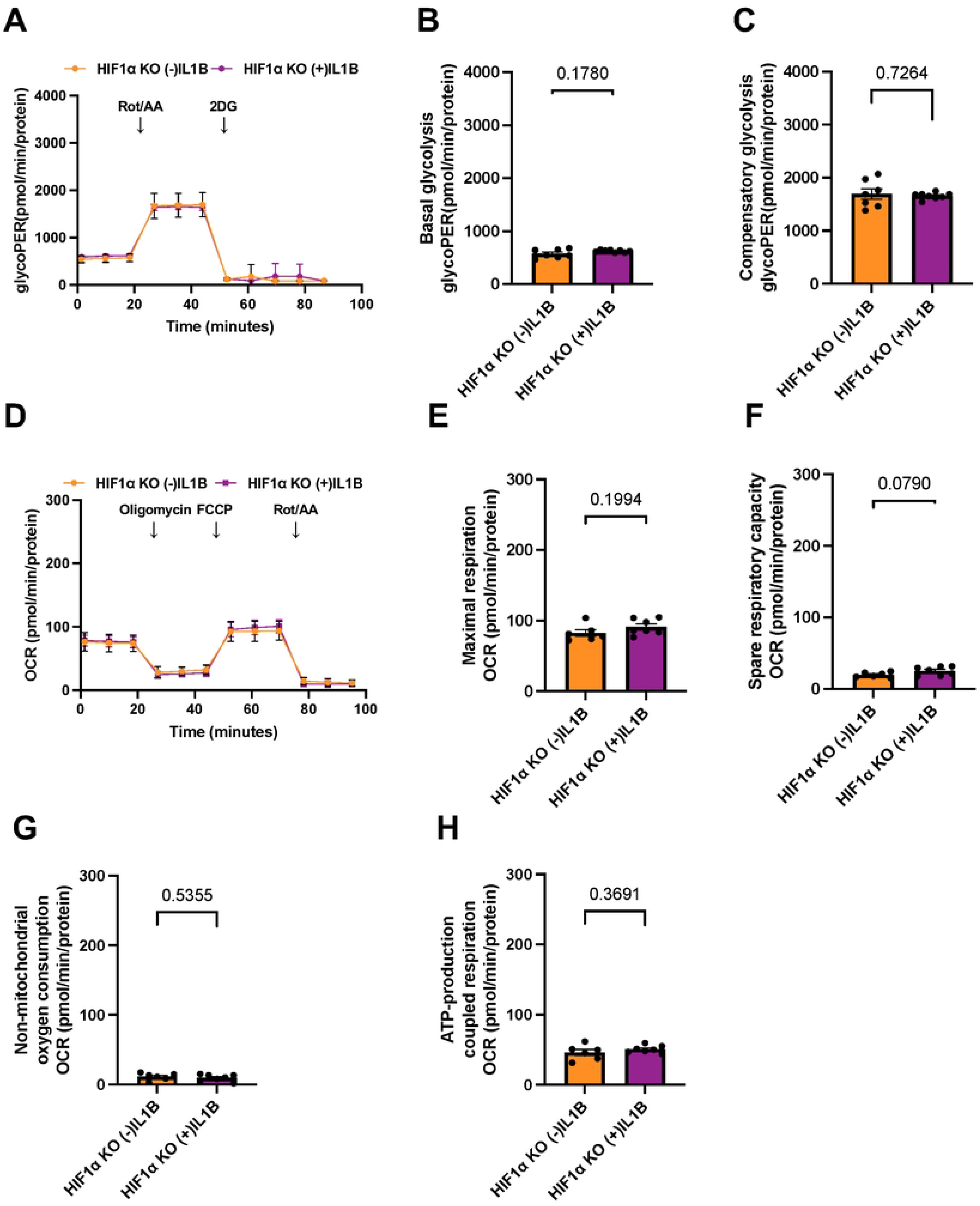
HIF1α knockout blocks interleukin-1β induced glycolysis. Contribution of glycolysis to proton efflux rate (glycoPER) (**A**), basal glycolysis (**B**), and compensatory glycolysis (**C**) in untreated and IL-1β-treated HCT 116 HIF1α knockout colorectal cancer cells. Oxygen consumption rate (OCR) relative to baseline in untreated and IL-1β-treated HCT 116 HIF1α knockout cells (**D**). The maximal mitochondrial respiration (**E**), spare respiratory capacity (**F**), non-mitochondrial oxygen consumption (**G**), and ATP-coupled respiration (**H**) in untreated or IL-1β-treated HIF1α knockout colorectal cancer cells. Each data point represents the average OCR over 3–5 replicates per condition. Error bars are mean ± SEM and ns indicates no statistical significance (P > 0.05).

### TNFα shares a similar mechanism utilizing AKT and HIF1α to increase glycolysis

Tumor necrosis factor alpha (TNFα), like IL-1β, has been reported to be elevated in ulcerative colitis and colorectal cancer [39–42]. Similar to IL-1β, we found that TNFα increased phospho-AKT and HIF1α levels in HCT 116 colorectal cancer cells (**Fig 6A**). Consistent with these effects, treatment of colorectal cancer cells with TNFα produced a significant elevation in the glycolytic Proton Efflux Rate (**Fig 6B**). Basal (**Fig 6C**) and compensatory (**Fig 6D**) glycolysis were higher in TNFα-treated cells. This raises the possibility that both IL-1β and TNFα could act through a similar mechanism to elevate glycolysis in these cells.

**Figure 6.**
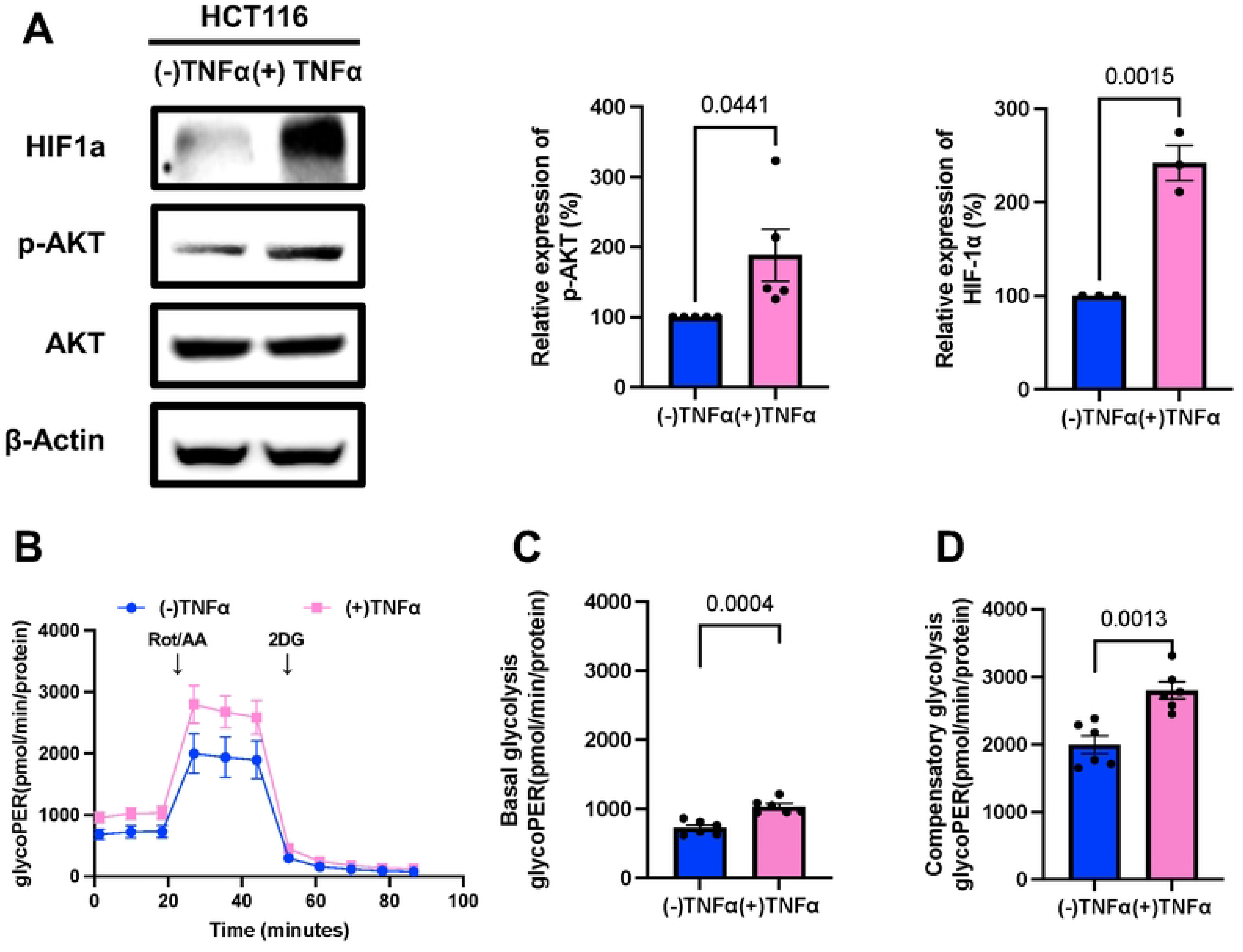
TNFα also increases HIF1α through AKT activation in colorectal cancer cells. (**A**) Western blot analysis of HIF1α and AKT expression in TNFα-treated or untreated HCT 116 cells with β-actin as a loading control. The right panels show the quantification of phospho-AKT (Ser473) relative to total AKT expression and HIF1α levels. Contribution of glycolysis to proton efflux rate (glycoPER) (**B**), basal glycolysis (**C**), and compensatory glycolysis (**D**) in untreated or TNFα-treated HCT 116 colorectal cancer cells. Each data point represents the average glycoPER over 3–5 replicates per condition. Error bars are mean ± SEM. Significant differences are shown.

### Increased glycolysis requires AKT and HIF1α in TNFα treated colorectal cancer cells

To test whether TNFα shares a similar mechanism, the effect of TNFα treatment in HCT 116 cells lacking HIF1α or AKT1/2 was assessed. TNFα-treated cells showed no change in glycolysis in HCT 116 cells with AKT1/2 knocked out (**Fig 7A**). Neither basal glycolysis (**Fig 7B**) or compensatory glycolysis (**Fig 7C**) was altered in these cells, thus demonstrating the requirement of AKT1/2 in mediating TNFα induction of glycolysis in colorectal cancer cells. HIF1α knockout produced a similar result, as these cells also failed to show a glycolytic response from TNFα treatment (**Fig 7D**). Basal (**Fig 7E**) and compensatory (**Fig 7F**) were the same regardless of TNFα treatment in colorectal cancer cells. These results confirm a shared mechanism between IL-1β and TNFα that involves activation of AKT and increased HIF1α.

**Figure 7.**
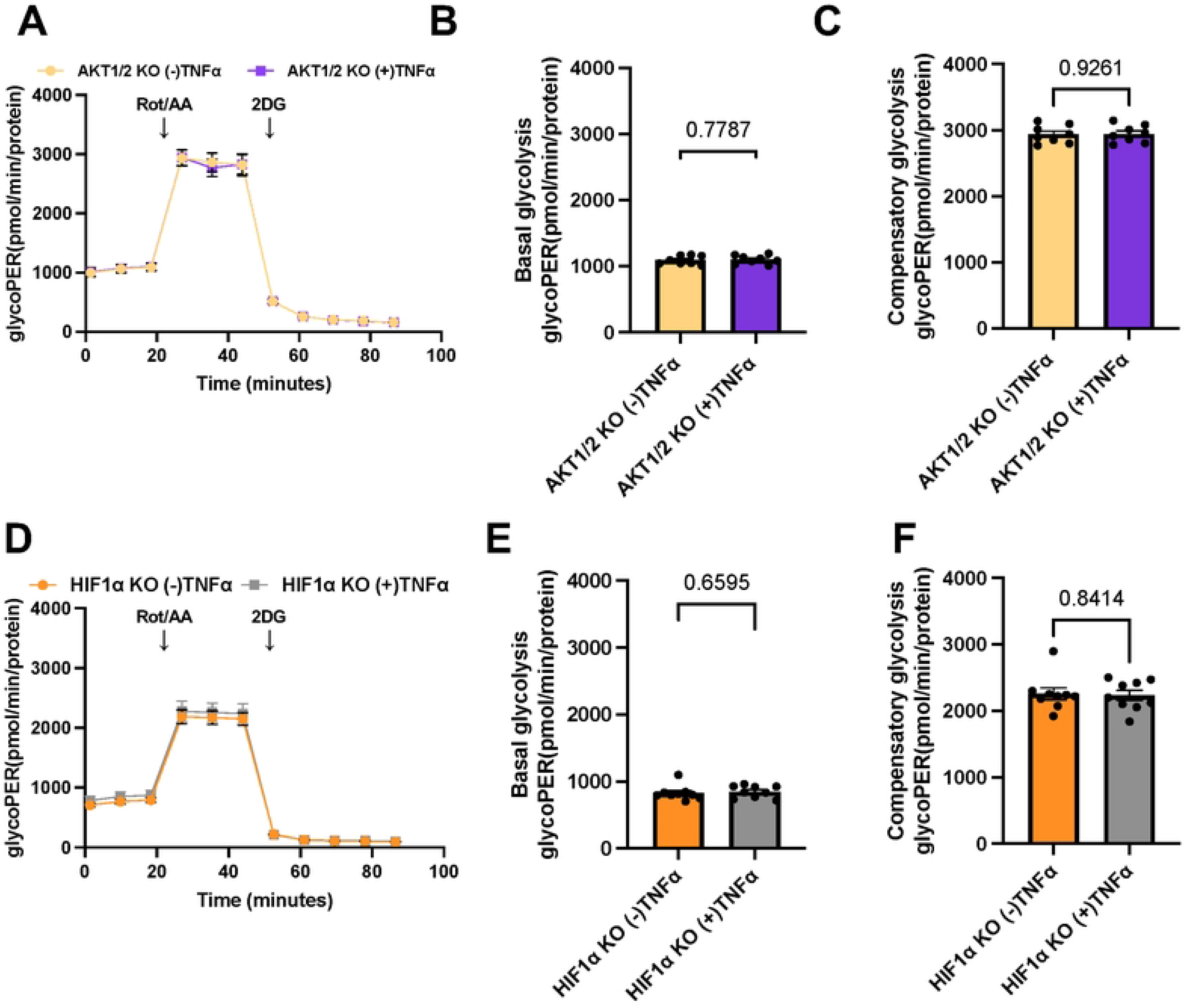
AKT1/2 and HIF1α knockout blocks TNFα-induced glycolysis. Contribution of glycolysis to proton efflux rate (glycoPER) (**A**), basal glycolysis (**B**), and compensatory glycolysis (**C**) in untreated or TNFα-treated AKT1/2 knockout HCT 116 colorectal cancer cells. Glycolytic component of the proton efflux rate (glycoPER) (**D**), basal glycolysis (**E**), and compensatory glycolysis (**F**) in un-treated or TNFα-treated HIF1α knockout colorectal cancer cells. Data points represent the average glycoPER over 3–5 replicates per condition. Error bars are mean ± SEM and ns indicates no statistical significance (P > 0.05).

## Discussion

Colorectal cancer cells predominantly rely on glycolysis to generate energy, even in the presence of oxygen (the Warburg effect), in contrast to non-cancerous colorectal cells, which primarily utilize butyrate oxidation [43]. This metabolic shift is influenced by several factors, including inflammation, specifically pro-inflammatory cytokines such as IL-1β and TNFα, which are expressed at higher concentrations in colorectal cancer [9, 44–46]. Proinflammatory cytokines also play a significant role in colorectal cancer development from initiating tumor initiation to progression [47]. In particular, genetic variations such as polymorphisms in the IL-1β gene have been associated with a higher likelihood of developing colorectal cancer [26, 48, 49]. Given the role of IL-1β in promoting glycolysis in colorectal cancer, we investigated its effects on colorectal cancer cell metabolism and found that IL-1β increases glycolysis through AKT and HIF1α in colorectal cancer cells.

Our findings expand previous observations on altered metabolism in colorectal cancer. Earlier work reported that colorectal cancer and ulcerative colitis have been shown to decrease the oxidation of butyrate and increase glycolysis [3, 18, 50]. Alterations in metabolic enzymes that regulate this change have been characterized; for example, our lab recently demonstrated that pyruvate kinase M1 (PKM1), which we found to be diminished in colorectal cancer samples compared to non-cancerous controls, promotes butyrate oxidation and suppresses glycolysis [3]. There are likely additional factors that also alter metabolism in colonic epithelial cells. Both colorectal cancer and ulcerative colitis are characterized by an elevated inflammatory state, where there are increases in several pro-inflammatory cytokines [36, 37, 51–53]. IL-1β is one of these cytokines that is significantly upregulated in ulcerative colitis and colorectal cancer, consistent with previous reports showing IL-1β promotes glycolysis in lung adenocarcinoma cells, macrophages, and colorectal cancer cells [19, 22, 23, 54]. Taken together, these findings highlight the importance of inflammatory cytokines in promoting altered metabolism.

Mechanistically, our data reveal that IL-1β-induced glycolysis requires activation of AKT and HIF1α. Consistent with our findings, IL-1β has been reported to activate HIF1α through an NFkB-dependent and cyclooxygenase-2 (COX-2)-dependent mechanism in A549 lung epithelial cancer cells [55]. AKT was identified as an upstream factor in HIF1α induction [56–58]. In another study, IL-1β and TNFα were shown to increase HIF1α activation or its binding to target genes in HepG2 cells [59]. Similarly, IL-1β has been reported to regulate the peptide adrenomedullin through a HIF1α mechanism in ovarian cancer cells and stimulate the production and secretion of stem cell factor in a PI3K/mTOR and HIF1α-dependent manner in leukemia cells [60, 61]. These findings suggest that this cytokine has a general role in regulating this transcription factor across diverse cell types. Interestingly, HIF1α has also been reported to mediate IL-1β transcription in macrophages [62].

In our study, HIF1α functions as a key downstream target that mediates the ability of this cytokine to drive metabolic changes, specifically increasing glycolytic flux. We were unable to determine how IL-1β elevates HIF1α through AKT, but mitochondrial dysfunction and increases in reactive oxygen species may represent a plausible mechanism that may be expanded to other cell types. Another possible mechanism is that IL-1β-induced AKT signaling could stabilize HIF1α through the activation of a mammalian target of rapamycin (mTOR) or inactivation of glycogen synthase kinase-3β (GSK3β) [56, 57]. In our experiments, we observed a significant increase in glycolysis after 6 h of IL-1β treatment, whereas oxidation of butyrate was suppressed only after 45 min of IL-1β treatment. This may suggest that the increased glycolysis observed through IL-1β treatment is a compensatory mechanism from the initial diminishment in butyrate oxidation. It would be interesting to test whether loss of butyrate oxidation, perhaps through knockout of short-chain acyl-dehydrogenase (ACADS), would block IL-1β’s ability to increase glycolysis.

Overall, our data presented here indicate that IL-1β and TNFα shift cellular metabolism toward glycolysis in cancerous colonocytes through activating AKT and increasing HIF1α. In our model, AKT is a downstream target that mediates elevated HIF1α levels, where both proteins are required for IL-1β or TNFα to promote glycolysis (**Fig 8**).

**Figure 8.**
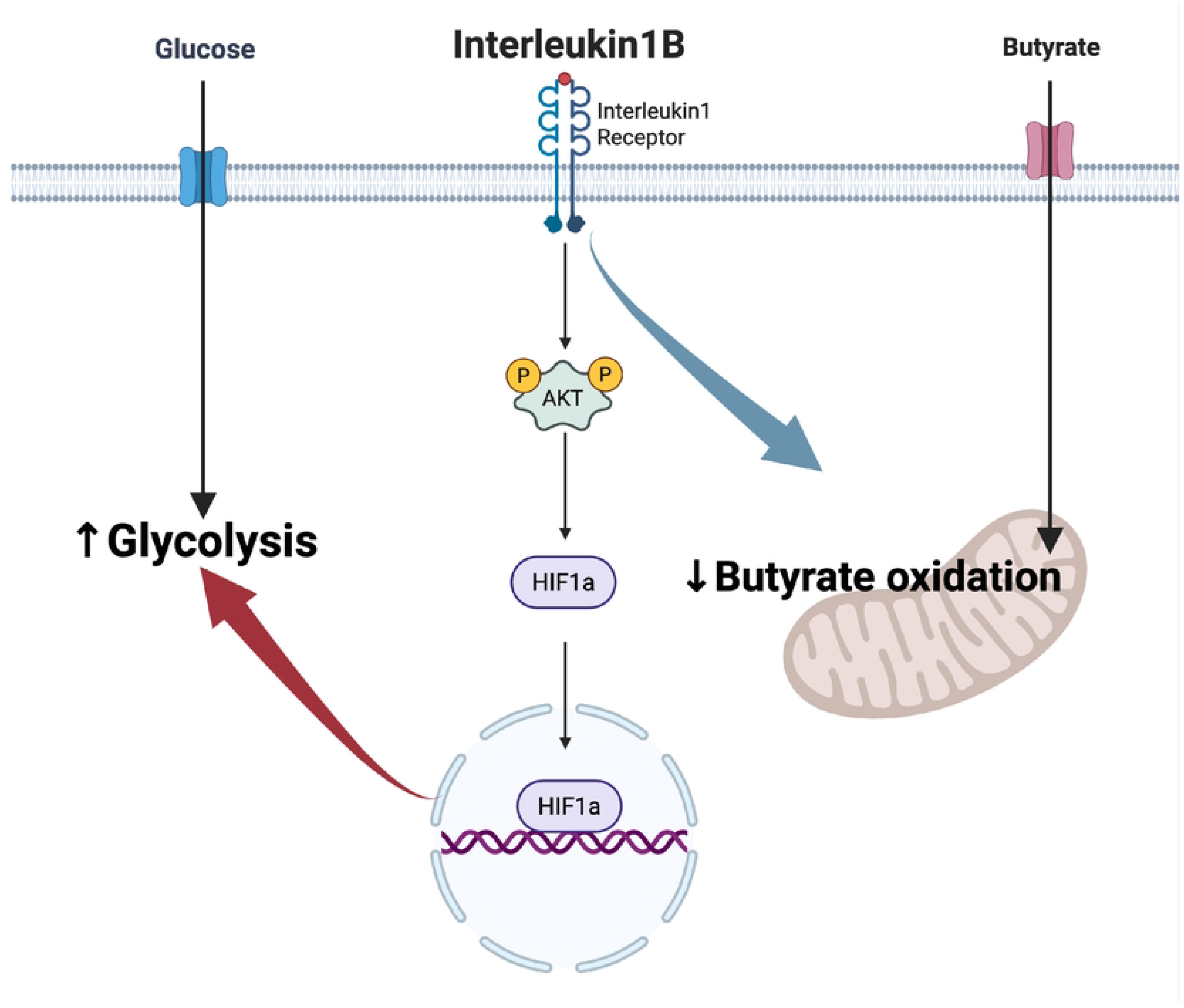
Mechanism of interleukin-1 beta induction of glycolysis in colorectal cancer cells. Experimental model of pathway involved in interleukin-1 beta upregulating glycolysis. In this model, the IL-1β binds to and signals through the interleukin-1 receptor. Activation of AKT increases HIF1α levels, which functions to shift cellular metabolism towards glycolysis. Created in BioRender. Kim, J. (2025) https://BioRender.com/1rsyzoy

A limitation of our study is that all experiments were conducted under normoxia. However, previous studies have shown that IL-1β induces HIF1α in several cell types, including human hepatocytes and breast cancer cells [63, 64]. Another report has demonstrated that TNFα enhanced HIF1α protein and mRNA levels via NF-kB pathway under both normoxia and hypoxia in human airway smooth muscle cells [65]. These findings support the physiological relevance of our results across normoxia and hypoxia. However, given that tumor microenvironment is generally hypoxic, future studies are needed to evaluate the effects of pro-inflammatory cytokines on metabolic shift under hypoxia or in the presence of cobalt chloride, which mimics hypoxia.

## Conclusions

In conclusion, there is a common mechanism between IL-1β and TNFα in mediating metabolic effects in colorectal cancer cells. Considering that both cytokines are elevated in the disease, this mechanism may help explain the role of these cytokines and inflammation in promoting cancer progression through increasing glycolysis in colorectal cancer cells.

## Author contributions

Conceptualization: J.K., D.R.D.; Methodology: J.K., B.P., A.B., D.R.D.; Validation: J.K., B.P., O.F.R., D.R.D.; Data Analysis: J.K., D.R.D.; Writing- Original Draft: J.K., D.R.D.; Writing- Editing and Review: J.K., A.B., D.R.D.; Supervision: D.R.D.; Project administration: D.R.D.; Funding acquisition: D.R.D.

## Acknowledgements

We would like to thank Dr. Bert Vogelstein and the Johns Hopkins Genetic Resources Core Facility for providing the parental HCT116, HIF1α knockout cell line (HIF1α ^-/-^), and AKT1/2 knockout cell line (AKT1/2^-/-^).

## Competing Interests

The authors declare that there are no conflicts of interest

## Funding

This work was supported by USDA NIFA (2019-67017-29261).

## Data Availability

All generated raw data and/or analyzed data from the current study are available from the corresponding author on reasonable request.

## Notes

### Competing Interest Statement

The authors have declared no competing interest.

